# Two Glu/Asp Residues Cooperatively Mediate an Early Step of ATP Hydrolysis in GHKL ATPases MutL and GyrB

**DOI:** 10.64898/2026.03.31.715682

**Authors:** Kenji Fukui, Ayaka Shibuya, Takeshi Murakawa, Takato Yano

**Affiliations:** Department of Biochemistry, Faculty of Medicine, Osaka Medical and Pharmaceutical University, 2-7 Daigaku-machi, Takatsuki, Osaka 569-8686, Japan; Department of Food Science and Nutrition, Faculty of Human Life and Environment, Nara Women’s University, Kitauoyanishi-machi, Nara, Nara 630-8506, Japan; Department of Chemistry, Faculty of Medicine, Osaka Medical and Pharmaceutical University, 2-7 Daigaku-machi, Takatsuki, Osaka 569-8686, Japan

## Abstract

GHKL ATPases share a unique Bergerat ATP-binding fold and regulate diverse biological processes through ATP-dependent conformational changes. An early step of ATP hydrolysis in this family has been attributed to a single highly conserved glutamate residue proposed to function as the general base. However, mutations of this residue impair both the ATPase activity and ATP binding, complicating interpretation of its catalytic role. Re-examination of the high-resolution crystal structures revealed a second conserved acidic residue positioned within a hydrogen-bonding distance from the nucleophilic water molecule. Using *Aquifex aeolicus* MutL and GyrB as model enzymes, we combined systematic mutagenesis, ATPase and ATP-binding assays, and X-ray crystallography to dissect the roles of these residues. We show that alignment of the nucleophilic water can be maintained as long as the conserved glutamate retains hydrogen-bonding capability, whereas efficient ATP hydrolysis requires proton-accepting capacity at least one of the two acidic residues. These results indicate that the conserved glutamate primarily governs positioning of the nucleophilic water, while activation of this water for catalysis is achieved through cooperative general base function of the two acidic residues. Extending this framework to human MutL homologs, PMS2 and MLH1, we showed that clinically reported variants of uncertain significance in these DNA mismatch repair proteins substantially reduced the ATPase activity, indicating functional impairment. Together, our findings refine the catalytic mechanism of GHKL ATPases and provide a structural and functional framework for interpreting disease-associated variants in GHKL ATPases. Phylogenetic and ancestral state analysis further indicated that the second acidic residue was likely to be present in the common ancestor of major GHKL ATPase lineages but was later modified in a branch including Hsp90, suggesting evolutionary remodeling of the catalytic mechanism in the branch.

## Introduction

The GHKL ATPase family comprises a wide range of proteins involved in diverse cellular processes, including DNA replication, DNA repair, protein homeostasis, chromatin regulation, and signal transduction ^1^. The GHKL ATPase family includes the following representative members: DNA gyrase B (GyrB), the ATPase subunit of type II topoisomerase that introduces negative supercoils into DNA ^2,3^; the molecular chaperone Hsp90, which regulates the folding and stability of numerous client proteins ^4^; histidine kinases of two-component regulatory systems, which function in environmental sensing and signal transduction ^5^; the DNA mismatch repair protein MutL, which maintains genome stability by correcting replication errors ^6,7^; and MORC family proteins, which contribute to chromatin organization and epigenetic regulation ^8,9^. Despite their functional diversity, these proteins share a conserved Bergerat ATP-binding motif and are thought to employ a common mechanism for ATP binding and hydrolysis ^1,10-13^.

For GHKL ATPases, ATPase activity is essential for protein function. In many cases, ATP binding and hydrolysis drive large-scale conformational changes that are coupled to biological activity. The ATP-bound and ADP-bound states often correspond to distinct oligomeric or domain arrangements, and cycling between these states enables GHKL ATPases to act as molecular switches or motors that coordinate complex cellular processes ^14-16^. A representative example of such functional coupling is provided by MutL, a key component of the DNA mismatch repair pathway ^17-19^. MutL consists of an N-terminal ATPase domain and a C-terminal nuclease domain, and ATP binding induces dimerization of the N-terminal domains, leading to rearrangement of the overall architecture and engagement of downstream repair factors ^14,20-22^. Subsequent ATP hydrolysis, stimulated by interacting proteins and DNA substrates, induces further conformational changes that activate its nuclease function and enable strand incision during mismatch repair ^7,23-25^. Thus, in MutL, the ATP binding and hydrolysis cycle directly controls large-scale structural transitions that are essential for its biological activity. A detailed understanding of the ATP hydrolysis mechanism is therefore crucial not only for elucidating the molecular basis of fundamental biological phenomena but also for clarifying disease mechanisms associated with dysfunction of GHKL ATPases.

In human, impaired function of MutL proteins compromises the DNA mismatch repair pathway, leading to reduced fidelity of DNA replication. Such defects are a major cause of Lynch syndrome, one of the most common hereditary cancer predisposition syndromes ^26-30^. Cancers associated with Lynch syndrome exhibit high responsiveness to immune checkpoint inhibitors ^31,32^, and genetic testing of MutL genes is recommended for diagnosis. However, clinical interpretation is often hampered by the presence of variants of uncertain significance. This limitation largely reflects the insufficient understanding of MutL function at the amino acid level. Variants of uncertain significance have also been identified in the ATPase active site of human MutL homologs, highlighting the importance of elucidating the ATP hydrolysis mechanism of MutL for establishing a molecular basis for genetic diagnosis of Lynch syndrome.

Among GHKL ATPases, GyrB, MutL, and MORC are thought to share particularly similar ATPase mechanisms ^11^. Within the Bergerat ATP-binding motif, a highly conserved glutamate residue, Glu29 in *Aquifex aeolicus* MutL (aqMutL), has been proposed to function as a general base that activates the nucleophilic water molecule for ATP hydrolysis ^33,34^. This proposal is based on observations that substitution of alanine or lysine for this glutamate abolishes the ATPase activity. In addition, a conserved lysine residue, Lys79, in aqMutL, has been suggested to act as a general acid, donating a proton to the reaction intermediate and thereby facilitating cleavage of the γ-phosphate ^11^. Other conserved basic residues have also been proposed to contribute as Lewis acids that stabilize negatively-charged reaction intermediates ^11,34^. Collectively, these residues have been proposed to form the core catalytic machinery of GHKL ATPases.

Previously, we determined high-resolution crystal structures of the ATPase domain of aqMutL in complex with ATP analogs, in which a water molecule presumed to be the nucleophile for ATP hydrolysis was clearly observed at a hydrogen-bonding distance to the conserved glutamate Glu29 ^35^. Upon careful re-examination of these structures, we noticed that this same water molecule is also positioned within a hydrogen-bonding distance to a second conserved acidic residue, Glu32 (Fig. 1A). Thus, the nucleophilic water is not exclusively coordinated by Glu29, but simultaneously interacts with two acidic residues.

**Figure 1.**
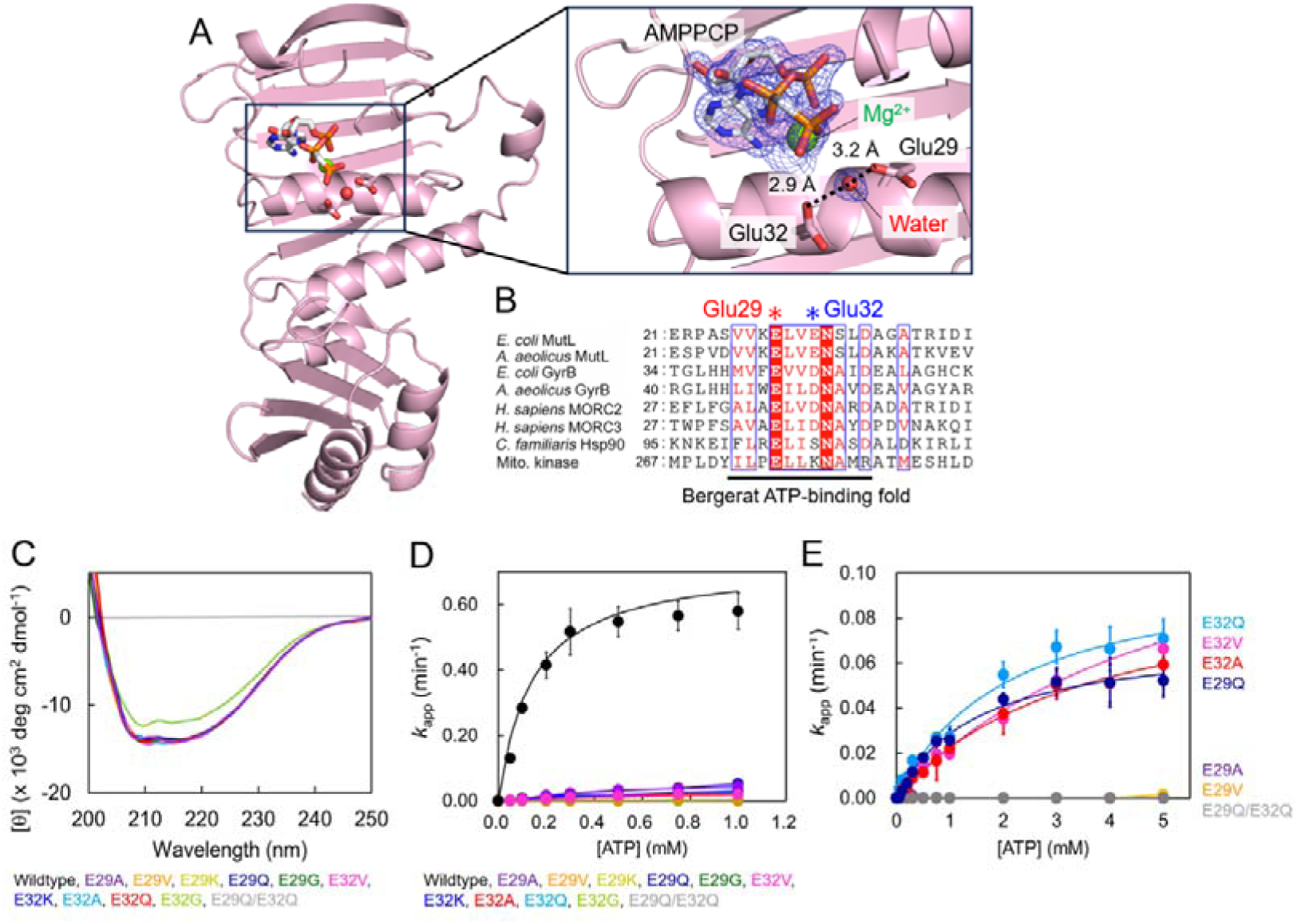
Effects of mutations on the overall structure and the ATPase activity of the aqMutL NTD. (A) Crystal structure of the aqMutL N-terminal domain (NTD) in complex with the non-hydrolyzable ATP analog AMPPCP. The boxed region indicates the ATP-binding site and is shown enlarged on the right. AMPPCP and the side chains of Glu29 and Glu32 are shown as stick models. Oxygen atoms are shown in red, nitrogen atoms in blue, and phosphorus atoms in orange. The magnesium ion and the water molecule putatively acting as the nucleophile are depicted as green and red spheres, respectively. Hydrogen bonds are indicated by dashed lines, with distances shown. The blue mesh represents an omit *F*_o_–*F*_c_ electron density map contoured at 3σ for AMPPNP, the magnesium ion, and the water molecule. (B) Sequence alignment of representative GHKL ATPases. The Bergerat ATP-binding fold is underlined, and the conserved acidic residues corresponding to aqMutL Glu29 and Glu32 are marked with asterisks. Amino acid sequence alignment was performed by CLUSTAL W program ^57^ and visualized by ESPript3 ^58^. (C) Far-UV CD spectra of the wildtype and mutant forms of the aqMutL NTD were collected and averaged over 10 consecutive scans. (D, E) ATPase activity of the aqMutL NTD was measured under steady-state conditions using a colorimetric assay that detects the release of inorganic phosphate. Apparent rate constants (*k*_app_) were calculated from the measured phosphate concentrations and plotted as a function of ATP concentration. Kinetic parameters were determined by fitting Michaelis–Menten equation to the data. Each data point represents the mean of three independent experiments, with error bars indicating standard deviation. Theoretical curves calculated from the fitted kinetic parameters are overlaid. Data for 0–1 mM substrate concentrations are shown to compare the wildtype from with mutant forms (D). Activities for higher substrate concentrations were measured for some of the mutant forms (E).

Although earlier biochemical studies, including those from our group, demonstrated that replacement of Glu29 with alanine abolishes the ATPase activity, such mutations also severely impair ATP binding ^11,33^, complicating interpretation of the specific catalytic role of this residue. In this context, the potential contribution of Glu32 to ATP hydrolysis might have been overlooked. Supporting this notion, the residue corresponding to Glu32 is conserved not only in MutL but also in other GHKL ATPases, including GyrB and MORC proteins (Fig. 1B). Furthermore, replacements of the corresponding Glu residue with Lys in *Escherichia coli* MutL and in human MutL homolog 1 (MLH1) have been reported to abolish DNA mismatch repair activity in cell-based or nuclear extract-based assays ^36,37^.

In this study, we investigated the functional role of Glu32 in the ATP hydrolysis mechanism of MutL and GyrB using a combination of enzymatic and X-ray crystallographic analyses. Our results provide new insights into the catalytic mechanism of GHKL ATPases and refine the current model of ATP hydrolysis for this protein family. By clarifying the molecular basis of ATP hydrolysis in GHKL ATPases, this work also provides a mechanistic framework for interpreting the functional consequences of some clinically observed variants, thereby contributing to the foundation for assessing the impact of variants of uncertain significance.

## Results and Discussion

### Mutational analysis of Glu29 and Glu32 in the ATPase domain of aqMutL

To examine the contribution of Glu32 to the ATPase activity of the aqMutL NTD and to directly compare its role with that of Glu29, we introduced a series of substitutions at both positions. To exclude the possibility that the observed effects on the ATPase activity arose from global structural perturbations, circular dichroism (CD) spectra were measured for all mutant forms (Fig. 1C). Among these, only the E32G mutant form exhibited a marked decrease in ellipticity, suggesting that this substitution disrupts the secondary structure of the protein. In contrast, all other mutant forms displayed CD spectra indistinguishable from that of the wildtype form, indicating that these substitutions do not substantially perturb the overall structure of the aqMutL NTD.

We first analyzed the effects of mutations at Glu29 on the ATPase activity of the aqMutL NTD. The mutation to Gln would selectively impair the proton association/dissociation capability of the Glu side chain while preserving its size and hydrogen-bonding potential. Moreover, because mutations to Val, Lys, and Gly at this position have been reported as variants of uncertain significance in human MutL, these mutations were also analyzed. Consistent with previous reports, all Glu29 mutant forms exhibited severe reduction in the ATPase activity (Fig. 1D and E, and Table 1). In particular, the E29A, E29V, E29K, and E29G mutant forms showed activities below the detection limit, indicating that Glu29 plays a central role in ATP hydrolysis. However, as described above, previous studies also demonstrated that mutations at Glu29 impair ATP binding ^11,33,34^, indicating that the loss of the ATPase activity of these mutant forms cannot be unambiguously attributed to disruption of a catalytic base function.

**Table 1.** Kinetic parameters for the ATPase activity of the aqMutL NTDs, aqGyrB NTDs, ProS2-tagged human PMS2 NTDs, and histidine-tagged human MLH1 NTDs.

| Proteins | $k_{\text{cat}}$ ($\text{min}^{-1}$)<sup>1</sup> | $K_{\text{m}}$ (mM)<sup>1</sup> | $k_{\text{cat}}/K_{\text{m}}$ ($\text{M}^{-1} \text{s}^{-1}$) |
| --- | --- | --- | --- |
| aqMutL WT | $0.65 \pm 0.10$ | $0.14 \pm 0.026$ | 77 |
| E29A | N.D.<sup>2</sup> | N.D.<sup>2</sup> | N.D.<sup>2</sup> |
| E29V | N.D.<sup>2</sup> | N.D.<sup>2</sup> | N.D.<sup>2</sup> |
| E29K | N.D.<sup>2</sup> | N.D.<sup>2</sup> | N.D.<sup>2</sup> |
| E29Q | $0.069 \pm 0.010$ | $1.5 \pm 0.20$ | 0.77 |
| E29G | N.D.<sup>2</sup> | N.D.<sup>2</sup> | N.D.<sup>2</sup> |
| E32A | $0.10 \pm 0.022$ | $2.1 \pm 0.39$ | 0.79 |
| E32V | $0.15 \pm 0.057$ | $5.5 \pm 1.3$ | 0.45 |
| E32K | N.D.<sup>2</sup> | N.D.<sup>2</sup> | N.D.<sup>2</sup> |
| E32Q | $0.10 \pm 0.018$ | $3.5 \pm 0.46$ | 0.48 |
| E32G | N.D.<sup>2</sup> | N.D.<sup>2</sup> | N.D.<sup>2</sup> |
| E29Q/E32Q | N.D.<sup>2</sup> | N.D.<sup>2</sup> | N.D.<sup>2</sup> |
| aqGyrB NTD WT | $2.4 \pm 0.19$ | $0.35 \pm 0.066$ | 110 |
| E48A | N.D.<sup>2</sup> | N.D.<sup>2</sup> | N.D.<sup>2</sup> |
| E48Q | $0.78 \pm 0.16$ | $0.75 \pm 0.34$ | 13 |
| D51A | $0.39 \pm 0.024$ | $0.21 \pm 0.038$ | 31 |
| D51N | $1.2 \pm 0.18$ | $0.28 \pm 0.059$ | 137 |
| E48Q/D51N | N.D.<sup>2</sup> | N.D.<sup>2</sup> | N.D.<sup>2</sup> |
| human PMS2 NTD WT | $0.62 \pm 0.049$ | $0.51 \pm 0.14$ | 20 |
| E44Q | $0.13 \pm 0.011$ | $0.22 \pm 0.079$ | 9.8 |
| E44V | $0.085 \pm 0.069$ | $0.67 \pm 0.14$ | 2.1 |
| human MLH1 NTD WT | $0.12 \pm 0.046$ | $0.66 \pm 0.064$ | 3.0 |
| E37K | N.D.<sup>2</sup> | N.D.<sup>2</sup> | N.D.<sup>2</sup> |

<sup>1</sup>The standard Michaelis-Menten equation was fitted to the data to determine the $k_{\text{cat}}$ and $K_{\text{m}}$ values.
<sup>2</sup>N.D. means that the activity was not detected.

To further dissect the role of Glu29, we focused on the E29Q mutant form. Using equilibrium dialysis, we confirmed that the E29Q mutant form binds ATP with an affinity comparable to that of the wildtype protein (Supplementary Fig. S1A). Because this substitution does not impair ATP binding, it should enable a more direct assessment of the proposed role of Glu29 as a general base in ATP hydrolysis. The E29Q mutant form retained measurable ATPase activity, although its *k*_cat_ value was approximately tenfold smaller than that of the wildtype enzyme (Fig. 1D and E, and Table 1). The persistence of substantial catalytic activity in the absence of the carboxylate group at the position 29 suggests that a residue other than Glu29 can participate in proton abstraction from the nucleophilic water during ATP hydrolysis.

We next examined the effects of mutations at Glu32. In contrast to the E29A mutant form of the aqMutL NTD, the E32A mutant form exhibited ATP binding ability comparable to that of the wildtype form (Supplementary Fig. S1A), indicating that Glu32 does not contribute to ATP binding. Mutations of Glu32 resulted in reduced ATPase activity relative to the wildtype form, supporting a role for this residue in catalysis. However, with the exception of the E32K and the structurally perturbed E32G mutant forms, all Glu32 mutant forms retained detectable ATPase activity (Fig. 1D and E, and Table 1), indicating that loss of Glu32 alone is insufficient to abolish ATP hydrolysis. Simultaneous substitutions of glutamine for the two residues in the E29Q/E32Q double mutant form led to a complete loss of the ATPase activity. Taken together, these results indicate that Glu29 and Glu32 do not exert equivalent functions but act cooperatively. The presence of at least one carboxylate group at either position, 29 or 32, appears to be required for ATP hydrolysis.

### Conserved roles of two acidic residues of aqGyrB in ATP hydrolysis

To examine whether the cooperative roles of Glu29 and Glu32 observed in aqMutL are conserved among other GHKL ATPases, we next analyzed the NTD of *A. aeolicus* GyrB (aqGyrB). A glutamate residue corresponding to aqMutL Glu29 is also conserved in bacterial GyrB and has been proposed to function as a general base in ATP hydrolysis ^38^. Although crystal structure of the *E. coli* GyrB NTD has been reported ^39^, the position of the putative nucleophilic water has not been clearly defined. Therefore, we selected the GyrB NTD from the hyperthermophilic bacterium *A. aeolicus* as a model system, anticipating that it would allow high-resolution structural analysis and clearer visualization of a catalytic water molecule.

First, we determined the crystal structure of this protein in complex with AMPPNP at 1.65 Å resolution (Table 2 and Fig. 2A). The overall structure was highly similar to that of the *E. coli* GyrB NTD, with a Cα root mean square deviation of 1.42 Å. As observed for the *E. coli* GyrB NTD, AMPPNP was accommodated by the Bergerat ATP-binding fold, and AMPPNP binding seemed to induce conformational changes that promoted dimer formation through the newly formed interface. Glu48 and Asp51 of aqGyrB correspond to Glu29 and Glu32 of aqMutL, respectively. In the crystal structure, these residues were found to tightly coordinate a water molecule that is likely to act as the nucleophilic water during ATP hydrolysis (Fig. 2A). Based on these structural observations, we introduced Ala or Gln/Asn substitutions for Glu48 and Asp51. The CD spectra of the mutant forms were nearly identical to that of the wildtype form (Fig. 2B), suggesting that these substitutions did not disrupt the overall structure of the aqGyrB NTD.

**Figure 2.**
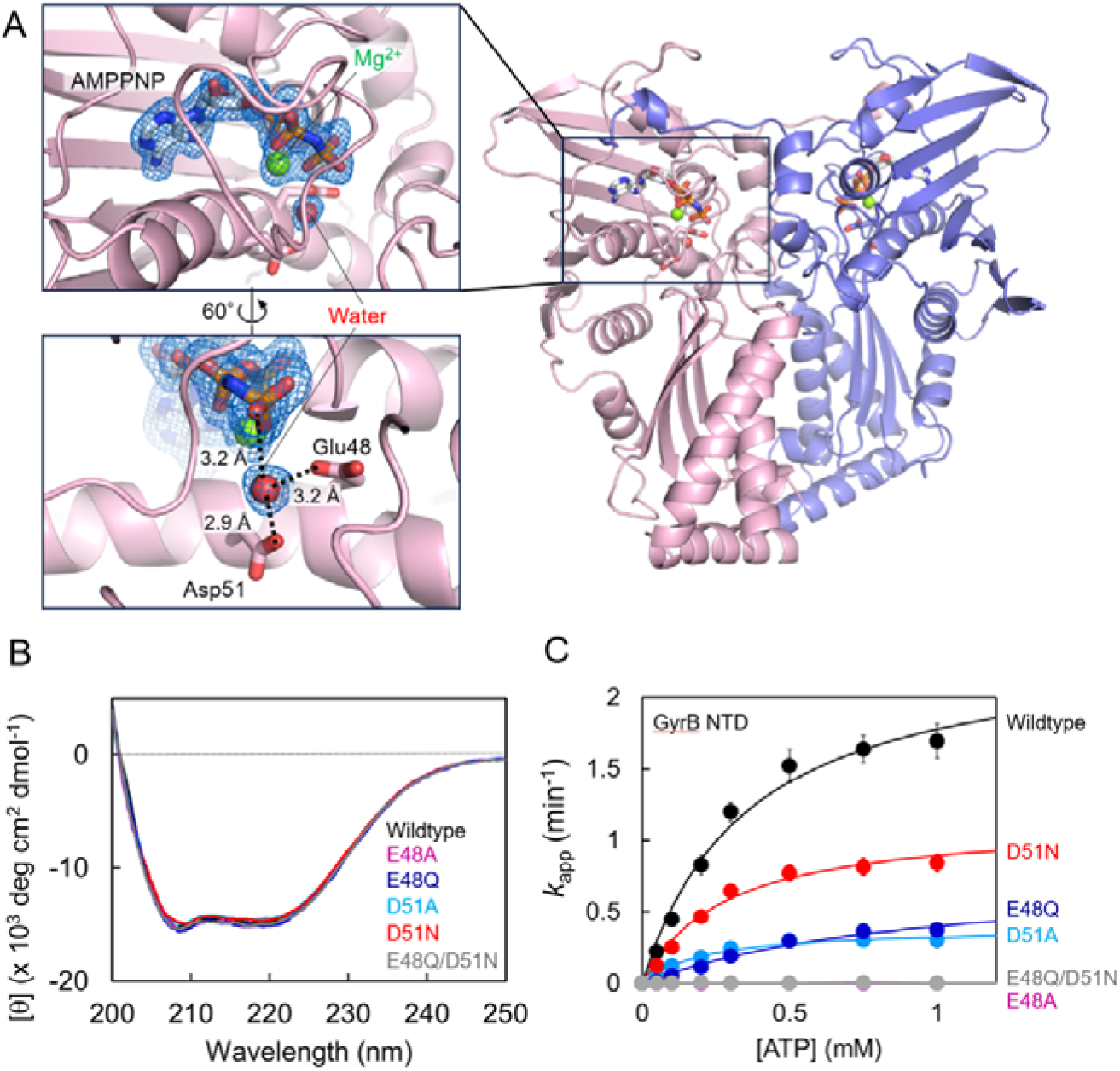
Crystal structure of the aqGyrB NTD and effects of mutations on the overall structure and the ATPase activity of the aqGyrB NTD. (A) Structure of the AMPPNP-bound dimer of the wildtype aqGyrB NTD. The boxed region highlights the ATP-binding site and is enlarged in the left panel. The same region is further enlarged and rotated 60° to provide an alternative view of the active site geometry. AMPPNP and the side chains of Glu48 and Asp51 are shown as stick models. Oxygen atoms are shown in red, nitrogen atoms in blue, and phosphorus atoms in orange. The magnesium ion and the water molecule presumed to act as the nucleophile are depicted as green and red spheres, respectively. Hydrogen bonds are indicated by dashed lines, with distances shown. The blue mesh represents an omit *F*_o_–*F*_c_ electron density map contoured at 3σ for AMPPNP, the magnesium ion, and the water molecule. (B) Far-UV CD spectra of the wildtype and mutant forms of the aqGyrB NTD were recorded and averaged over 10 accumulations. (C) ATPase activity of the aqGyrB NTD was measured under steady-state conditions as described in Fig. 1. The *k*_app_ values were plotted as a function of ATP concentration. Michaelis–Menten equation was fitted to the data. Data points represent the mean ± standard deviations from three independent experiments.

**Table 2.**
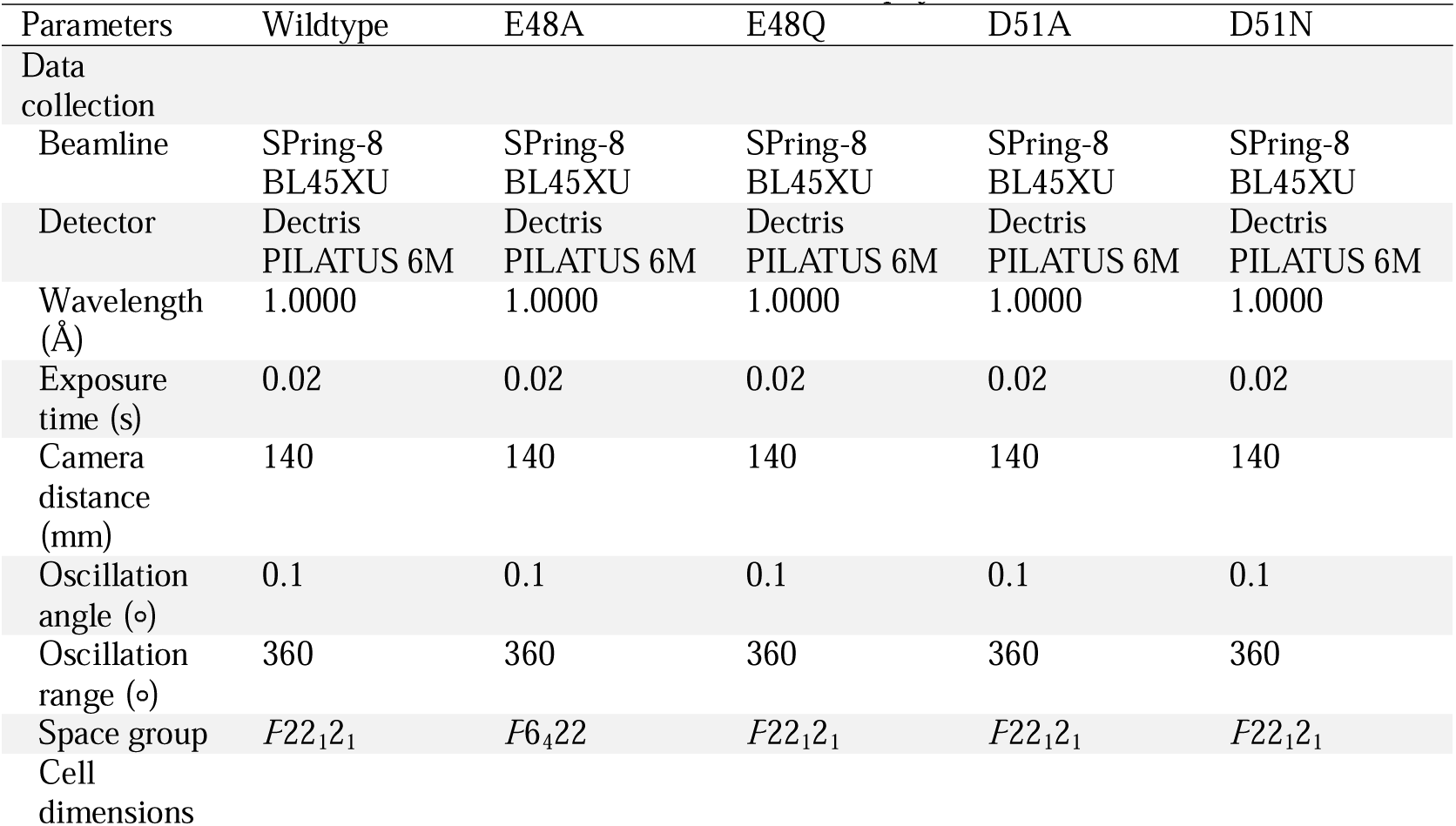
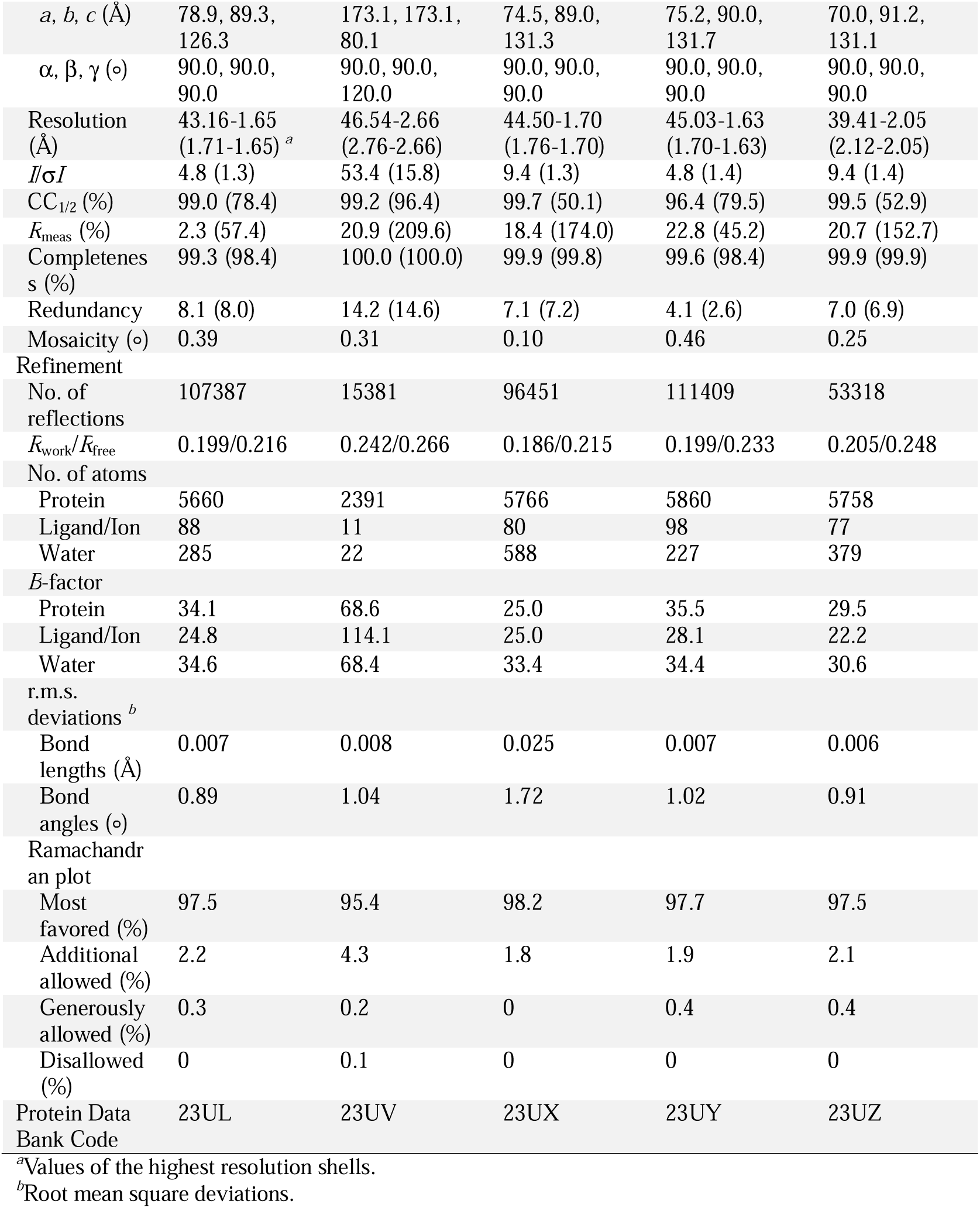
Data collection and refinement statistics for the aqGyrB NTDs.

Equilibrium dialysis showed that the E48A mutant form failed to bind ATP, whereas the E48Q mutant form displayed ATP-binding affinity comparable to that of the wildtype protein (Supplementary Fig. S1B). The E48Q mutation therefore allowed us to evaluate the catalytic contribution of Glu48 independently of ATP binding. ATPase activity assays showed that the E48A mutant form completely lacked the activity, while the E48Q mutant form retained approximately one third of the wildtype activity (Fig. 2C and Table 1). The persistence of substantial catalytic activity in the E48Q mutant form suggests that another residue partially compensates for the loss of the Glu48 carboxylate in the proton abstraction.

The D51A mutant form of the aqGyrB NTD retained ATP binding ability comparable to that of the wildtype form, indicating that Asp51 is not required for nucleotide binding (Supplementary Fig. S1B). Mutations of Asp51 also impaired ATP hydrolysis, with the D51A and D51N mutant forms retaining ∼16% and ∼50% of the wildtype activity, respectively (Fig. 2C and Table 1). Notably, the E48Q/D51N double mutant form exhibited no detectable ATPase activity, resembling the effect of the corresponding double mutation in the aqMutL NTD. Taken together, these results indicate that both Glu48 and Asp51 contribute to the catalytic mechanism of ATP hydrolysis in aqGyrB and that at least one of these residues must retain a carboxylate group with the proton-accepting capability to sustain the enzymatic activity.

### Crystal structures of aqGyrB mutant forms and the role of Glu48 in ATP binding

In order to relate the ATPase activity profiles of the mutant forms to their structural features, we determined X-ray crystal structures of selected mutant forms using the aqGyrB NTD as a model system, which allows relatively high-resolution structural analyses. The AMPPNP-bound structures of the E48A, E48Q, D51A, and D51N mutant forms were determined at resolutions of 2.66, 1.70, 1.63, and 2.05 Å, respectively (Table 2). To show that the introduced substitutions were unambiguously supported by the crystallographic data, electron density maps around residues 48 and 51 are shown in Supplementary Fig. S2.

Structural superposition of the aqGyrB NTD and aqMutL NTD revealed that the catalytic Mg² ion occupies essentially the same position in the two ATPase active sites (Supplementary Fig. S3), indicating that the metal-binding geometry is highly conserved, where the Mg^2+^ ion is coordinated by the side chain of Asn52, AMPPNP, and surrounding water molecules. Neither Glu48 nor Asp51 directly coordinated the Mg^2+^ ion.

Since the conserved general base Glu (i.e. Glu48 of aqGyrB) does not directly interact with ATP, it has been unclear why mutations at the Glu residue abolish the ATP-binding ability of GHKL ATPases. To clarify this, we compared the crystal structures of the wildtype and mutant forms of the aqGyrB NTD (Fig. 3). In GHKL ATPases, ATP binding is tightly coupled to conformational rearrangements, including formation of an ATP lid (Fig. 3A), which acts to secure the bound nucleotide by enclosing it within the active site ^3,21^. Structural analysis of the wildtype aqGyrB NTD revealed that the side chain of Glu48 forms a hydrogen bond with Gln340, thereby stabilizing a loop containing Gln340 in a conformation capable of ATP binding (Fig. 3B). In this arrangement, Gln340 in turn forms a hydrogen bond with His122, a residue located within the ATP lid, while Lys342 in the same loop is positioned to interact with the γ-phosphate of ATP. These coordinated interactions appear to promote the ATP-induced conformational transition required for stable nucleotide binding.

**Figure 3.**
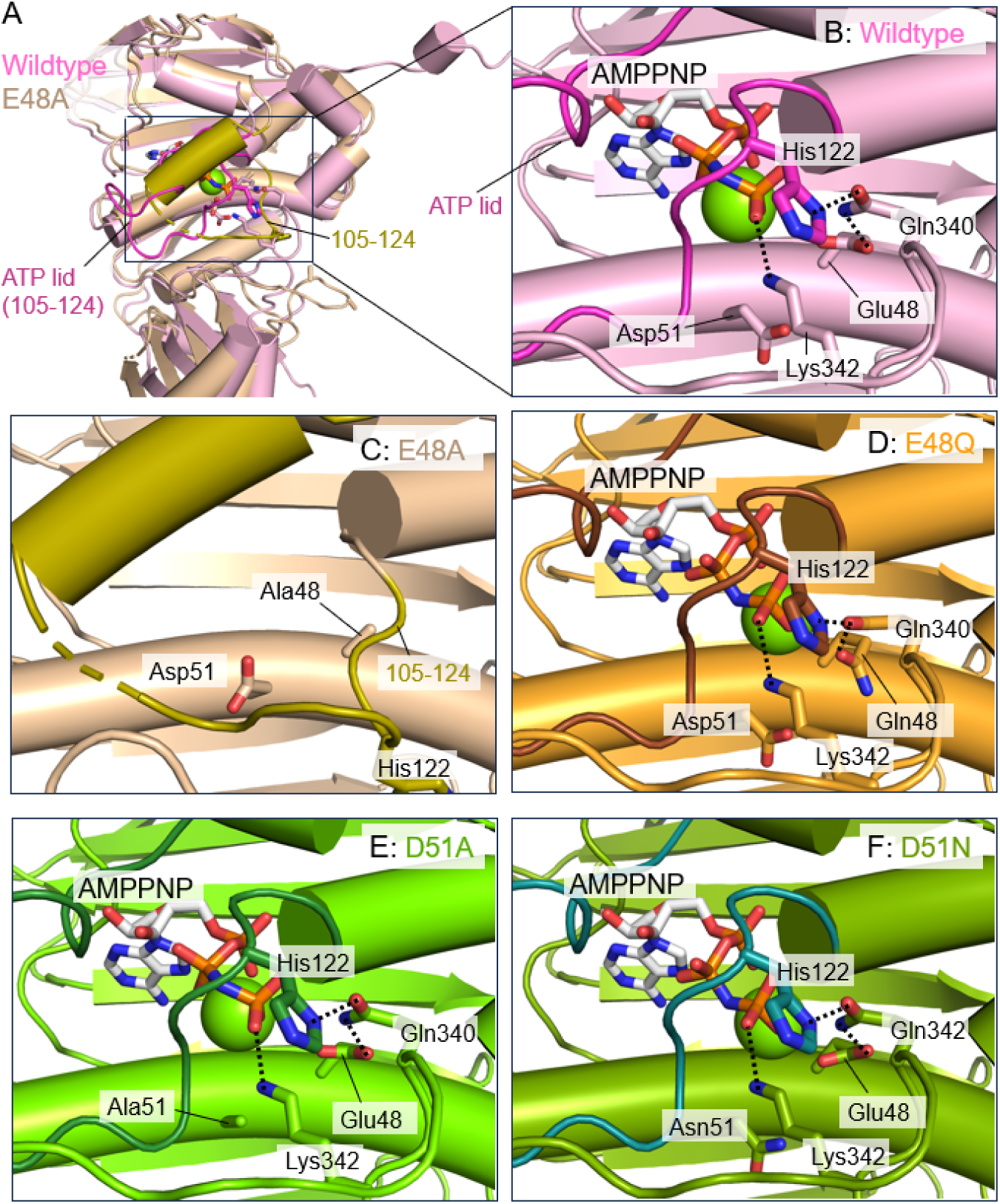
Structural basis for the loss and preservation of ATP binding in aqGyrB NTD mutant forms. The wildtype and all mutant forms were crystallized in the presence of AMPPNP; however, the E48A mutant form did not bind AMPPNP, whereas the E48Q, D51A, and D51N mutant forms were observed in complex with AMPPNP. (A) Overall structures of the wildtype and E48A mutant aqGyrB NTD monomers. The ATPase active site is indicated by a boxed region. The ATP-lid (residues 105–124) in the wildtype form and the corresponding region in the E48A mutant form are shown in magenta and olive, respectively. (B–F) Enlarged views of the ATPase active site in the wildtype (B), E48A (C), E48Q (D), D51A (E), and D51N (F) forms. Side chains of the residues implicated in the ATP-induced conformational changes (e.g. His122, Gln340, and Lys342), as well as residues 48 and 51, are shown as stick models. AMPPNP is depicted as sticks, and the Mg² ion is shown as a green sphere. Hydrogen bonds and ionic interactions are indicated by dashed lines.

In contrast, structural analysis of the E48A mutant form demonstrated that disruption of the interaction with Gln340 abolishes this network of hydrogen bonds, preventing the associated conformational rearrangements and thereby impairing ATP binding (Fig. 3C). These findings indicate that the interaction network emanating from Glu48 is indispensable for the conformational changes associated with the ATP lid formation. Interestingly, in the E48Q mutant form, the hydrogen bond with Gln340 is preserved, allowing the conformational transition to occur and enabling ATP binding (Fig. 3D). Consistently, in the D51A and D51N mutant forms, the interaction mediated by Glu48 remains intact, and the structural rearrangements required for ATP binding are maintained (Fig. 3E and F).

These structural observations extend previous models of GyrB ATPase function by linking the conserved catalytic glutamate to ATP lid stabilization and nucleotide-dependent conformational rearrangement. The E48A aqGyrB NTD structure, in particular, provides direct structural evidence that disruption of this Glu-centered interaction network is sufficient to prevent ATP-induced active-site assembly. This hydrogen-bonding network emanating from the general base Glu is conserved in GyrBs from other organisms ^39,40^ (Supplementary Fig. S4), while it is not observed in other GHKL ATPases, including MutL ^22,34,35,41^ and MORC family proteins ^9,42,43^. Therefore, the loss of ATP-binding ability observed upon the mutation of Glu29 to Ala in MutL ^11^ is likely to be explained by a different mechanism. This difference highlights mechanistic diversity within the GHKL ATPase family, despite conservation of the two catalytic acidic residues.

### Nucleophilic water molecule in the active site of mutant forms of the aqGyrB NTD

Having established the structural basis for the altered ATP-binding properties of the mutant forms, we next analyzed how these substitutions influence the positioning of the nucleophilic water molecule. Structural comparisons of the catalytic sites of the four mutant forms are shown in Fig. 4. In the E48A mutant form (Fig. 4A), the nucleophilic water molecule was not observed in the catalytic site. In contrast, in the E48Q mutant form, the nucleophilic water molecule was positioned at essentially the same location as in the wildtype form, coordinated by Gln48 and Asp51, and bound AMPPNP was observed (Fig. 4B). This structural arrangement is consistent with the biochemical data described above, in which the E48Q mutant form retained substantial ATP-binding and ATPase activities.

**Figure 4.**
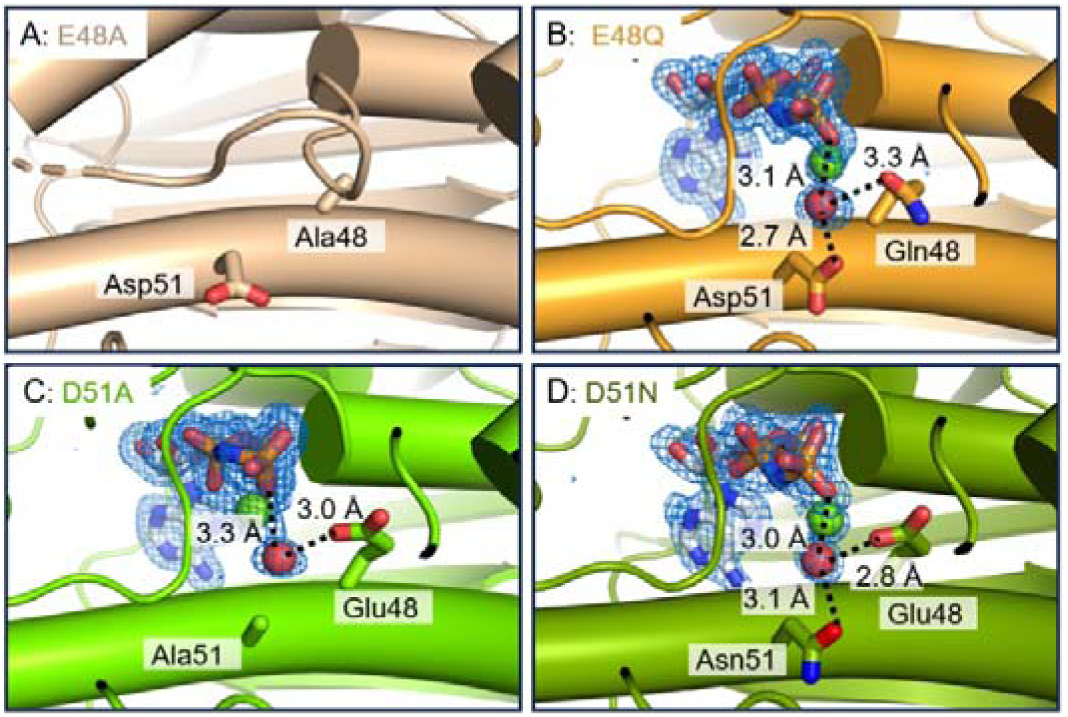
Structures surrounding the conserved acidic residues: (A) E48A, (B) E48Q, (C) D51A, and (D) D51N mutant forms of the aqGyrB NTD. AMPPNP and the side chains at the positions 48 and 51 are shown as stick models. Oxygen atoms are shown in red, nitrogen atoms in blue, and phosphorus atoms in orange. The magnesium ion is shown as a green sphere, and the water molecule proposed to act as the nucleophilic water is shown as a red sphere. Blue mesh represents *F*_o_–*F*_c_ omit electron density maps contoured at 3σ, calculated after omission of AMPPNP, the magnesium ion, and the nucleophilic water molecule. Hydrogen bonds are indicated by dashed lines, with the corresponding distances shown.

Unlike the E48A mutant form, the D51A mutant form bound AMPPNP (Fig. 4C), and the nucleophilic water molecule was coordinated only by Glu48. This asymmetric coordination suggests that Glu48 plays a dominant role in positioning the nucleophilic water molecule. In the D51N mutant form (Fig. 4D), which lacks the proton-transfer capability of Asp51 while retaining its hydrogen-bonding capacity, the nucleophilic water was coordinated by both Glu48 and Asn51. These structural features are consistent with the relative ATPase activities observed for the mutant forms of these two acidic residues (Fig. 2C).

Taken together, these results indicate that Glu48 and Asp51 make distinct yet cooperative contributions to ATP hydrolysis. Glu48 appears to play a dominant role in aligning the nucleophilic water molecule and activating this water molecule through proton abstraction as a general base. In contrast, Asp51 seems to contribute less to the precise positioning of the nucleophilic water but participates cooperatively with Glu48 in the proton abstraction, thereby supporting efficient ATP hydrolysis. Although mutation of the two acidic residues impaired ATPase activity in both aqMutL and aqGyrB, the magnitude of the effects differed substantially, with a much greater reduction in catalytic efficiency in aqMutL than in aqGyrB. The molecular basis for this quantitative difference is currently unclear. One possible explanation is that subtle differences in the active-site architecture and surrounding residues alter the relative contribution of each acidic residue to catalysis, allowing aqGyrB to tolerate perturbation of either residue more effectively than aqMutL.

Notably, an acidic residue corresponding to Asp51 of aqGyrB is conserved among several GHKL ATPases, including MutL, GyrB, and MORC family proteins (Fig. 1B), suggesting that the two-acidic-residue mechanism proposed here may be shared among these subgroups. The second acidic residue is not conserved in other members of the GHKL family, such as Hsp90 and mitochondrial histidine kinases (Fig. 1B), where a single acidic residue is thought to fully function as the general base responsible for water activation. A similar divergence appears to apply to the previously-reported general acid Lys residue ^11^, further supporting the notion that GHKL ATPases can be divided into at least two mechanistically distinct subclasses. This mechanistic bifurcation provides insights into the molecular evolution of the GHKL ATPase family.

### Functional and clinical implications of the second acidic residue in human MutL homologs

Our analyses of the aqMutL NTD and aqGyrB NTD indicate that, in addition to the Glu residue traditionally considered to function as the general base, an additional conserved acidic residue (Glu or Asp) also participates in the general base catalysis. In human MutL homologs, this residue corresponds to Glu44 in Postmeiotic Segregation Increased 2 (PMS2) and Glu37 in MLH1. Notably, variants of uncertain clinical significance, including E44V (ClinVar variation ID: 232732), E44G (230139), and E44Q (186069) in PMS2, as well as E37G (230139) and E37K (89640) in MLH1, have been reported at these positions.

To examine whether these residues similarly contribute to the ATPase activity in human MutL homologs, we introduced the corresponding mutations into the NTDs of human PMS2 and MLH1 and measured their ATPase activities. The recombinant E44G human PMS2 NTD and E37G human MLH1 NTD could not be expressed successfully, which is consistent with our finding that the corresponding mutation in the aqMutL NTD disrupted the structural integrity of the protein (Fig. 1C). Both the E44V and E44Q mutant forms of the PMS2 NTD exhibited a marked reduction in the ATPase activity, retaining approximately one-sixth of the wildtype activity (Fig. 5A and Table 1). In contrast, the E37K mutation in the MLH1 NTD completely abolished the ATPase activity under our assay conditions (Fig. 5B and Table 1) unlike the corresponding glutamine substitutions, which retained substantial residual ATPase activity in the aqMutL, aqGyrB, and PMS2 NTDs. A similar complete loss of ATPase activity was also observed for the E32K mutant form of the aqMutL NTD. These observations suggest that the severe defect caused by the lysine substitution cannot be attributed simply to loss of the catalytic carboxylate. Instead, introduction of a positively charged side chain (charge reversal) is likely to perturb the local electrostatic environment. Structural characterization of the MLH1 E37K and aqMutL E32K mutant forms will be required to clarify the molecular basis of this severe functional defect.

**Figure 5.**
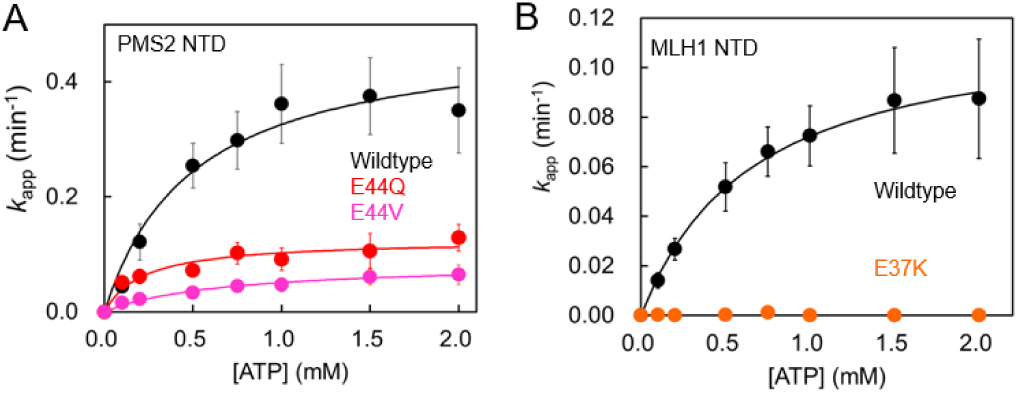
ATPase activities of human MutL homolog NTDs. (A) ATPase activity of mutant forms of the human PMS2 NTD. (B) ATPase activity of a mutant form of the human MLH1 NTD. ATPase activities were measured using the same procedures as described for Fig. 1 and Fig. 2. Apparent rate constants (*k*_app_) were plotted as a function of substrate concentration. Data points represent the mean values from three independent measurements, with error bars indicating standard deviations. Solid lines represent the theoretical Michaelis-Menten curves.

These results are in good agreement with our observations for the aqMutL NTD and aqGyrB NTD, further supporting the notion that this second acidic residue plays a conserved and functionally significant role in ATP hydrolysis among some members of the GHKL ATPase family. Previous studies demonstrated that even a several-fold reduction in the MutL ATPase activity could severely compromise mismatch repair efficiency *in vivo* ^11^. In this context, the substantial reduction in the ATPase activity caused by the E44V and E44Q mutations in the PMS2 NTD is likely to impair mismatch repair in human cells. In individuals carrying these germline variants, subsequent loss or inactivation of the remaining wildtype allele would leave only the ATPase-defective MutL protein, thereby compromising mismatch repair and promoting tumorigenesis. Nevertheless, whether these variants indeed compromise mismatch repair activity *in vivo* remains to be directly tested. Functional validation using cell-based or nuclear extract-based mismatch repair assays, such as the 6-thioguanine sensitivity assay ^44^ or CIMRA assay ^36^, will be required to establish a definitive link between the reduced ATPase activity and mismatch repair deficiency in human cells.

Although we focused here on pathogenic variants in the MutL homologs MLH1 and PMS2, extending similar structural and biochemical analyses to disease-associated variants in other human GHKL ATPases will be important for evaluating the generality and clinical relevance of the conserved catalytic mechanism proposed in this study.

### Phylogenetic analysis of GHKL ATPases

The newly-identified functionally-important acidic residue is not uniformly conserved across the GHKL superfamily. We therefore used this position as an evolutionary marker to investigate patterns of molecular divergence within the GHKL ATPase domain. In histidine kinases, even the canonical catalytic glutamate is typically replaced, and the ATPase-associated catalytic machinery is not maintained ^1,45,46^. Therefore, phylogenetic reconstruction was restricted to other GHKL ATPase family members MutL, GyrB, MORC, and Hsp90.

Phylogenetic analysis based on the trimmed ATPase-domain alignment recovered these major GHKL ATPase lineages (MutL, GyrB, MORC, and Hsp90) as a well-supported cluster, indicating relatively close evolutionary relationships among these groups, which is consistent with a previous study ^1^ (Supplementary Fig. S5). Ancestral state reconstruction performed on the maximum-likelihood phylogeny strongly supports the presence of an acidic residue at the position corresponding to aqMutL Glu32 or aqGyrB Asp51 in the internal node uniting the MutL, GyrB, MORC, and Hsp90 clades with a posterior probability of 0.983 (Fig. 6). In contrast, the ancestral node of the Hsp90 clade is strongly supported to encode serine at this position with a posterior probability of 1.000. Because no external outgroup was specified, the global polarity of amino acid residue changes across the entire phylogeny cannot be formally determined. Nevertheless, the widespread conservation of the second acidic residue across the MutL, GyrB, and MORC lineages, together with the high posterior probability at the most recent common ancestor of these groups and Hsp90, indicates that this residue was very likely to be present in their ancestor. Its absence within the Hsp90 lineage is therefore most parsimoniously interpreted as lineage-specific loss. In summary, these results suggest that the Hsp90 lineage underwent remodeling of its ATPase catalytic mechanism during its evolutionary divergence from other GHKL ATPase family members.

**Figure 6.**
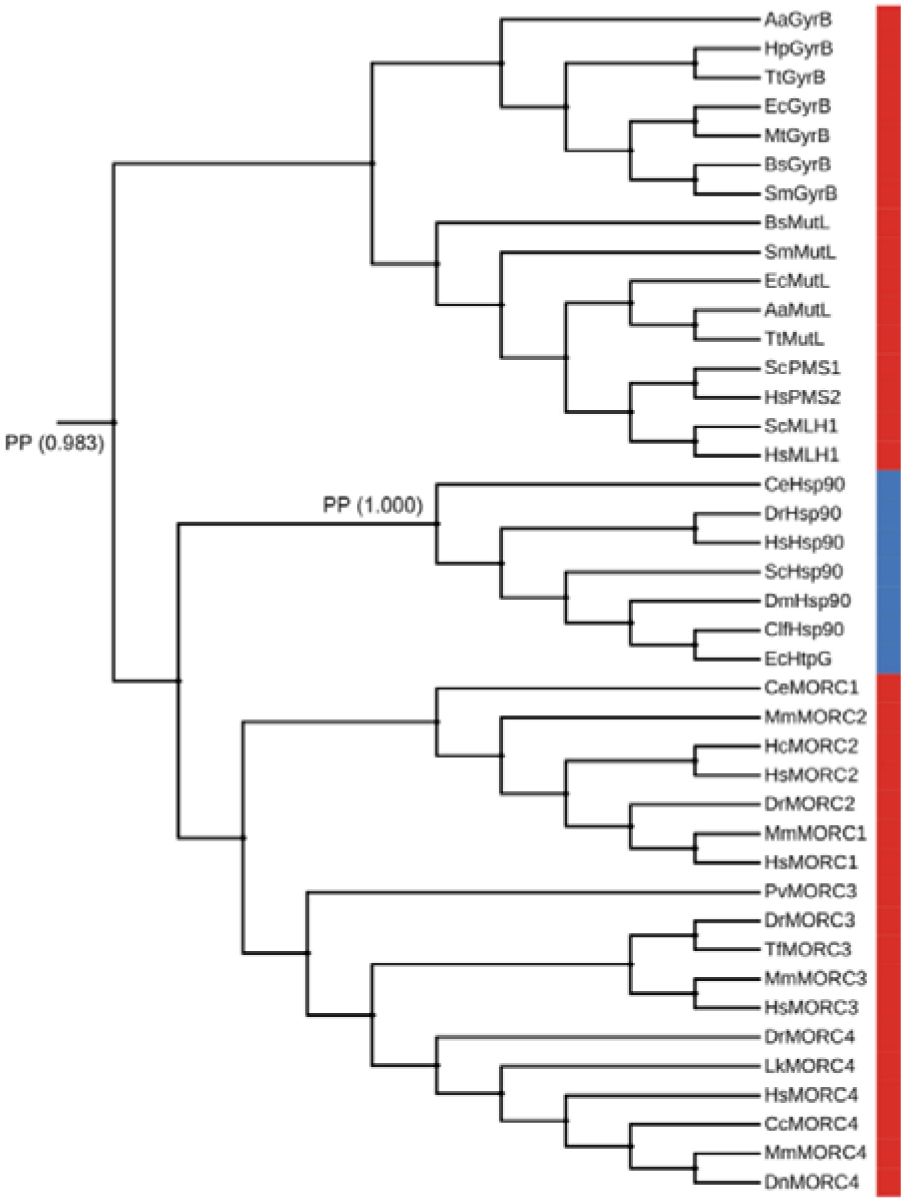
Maximum-likelihood phylogeny of the ATPase domain of representative GHKL family members. The state of the amino acid residue at the position corresponding to aqMutL Glu32 or aqGyrB Asp51 is indicated by colored bars (red, acidic residue; blue, non-acidic residue). The phylogenetic tree was visualized and annotated using the Interactive Tree of Life (iTOL) web server ^56^. The tree is displayed in a rectangular layout for clarity and should be interpreted as unrooted. Ancestral state reconstruction was performed on the fixed topology, and posterior probabilities are shown for selected internal nodes. The internal node uniting the MutL/GyrB/MORC/Hsp90 clades strongly supports an acidic residue with the posterior probability (PP) of 0.983, whereas the Hsp90 stem ancestor is strongly supported to encode serine at this position with the posterior probability of 1.000. Species abbreviations used in the phylogenetic tree are as follows: Ce, *Caenorhabditis elegans*; Sc, *Saccharomyces cerevisiae*; Dr, *Danio rerio*; Hs, *Homo sapiens*; Aa, *Aquifex aeolicus*; Hp, *Helicobacter pylori*; Tt, *Thermus thermophilus*; Ec, *Escherichia coli*; Mt, *Mycobacterium tuberculosis*; Bs, *Bacillus subtilis*; Sm, *Streptococcus mutans*; Dm, *Drosophila melanogaster*; Clf, *Canis lupus familiaris*; Mm, *Mus musculus*; Hc, *Hemicordylus capensis*; Pv, *Patella vulgata*; Tf, *Tachysurus fulvidraco*; Lk, *Lepidochelys kempii*; Cc, *Castor canadensis*; Dn, *Dasypus novemcinctus*.

## Conclusions

In this study, we re-examined the catalytic mechanism of GHKL ATPases using aqMutL and aqGyrB as model systems with mutational, biochemical, and crystallographic studies. Our results reveal that ATP hydrolysis in these enzymes is not governed by a single general base residue, as has been traditionally assumed, but instead relies on the coordinated action of two conserved acidic residues. Structural analyses indicate that Glu48 of aqGyrB plays a dominant role in aligning the nucleophilic water molecule in an optimal geometry for attack on the γ-phosphate of ATP. However, biochemical and structural evidence from single and double mutant forms indicates that both residues contribute to proton abstraction from the nucleophilic water. Thus, general base catalysis in GyrB and MutL appears to be achieved by redundant proton-accepting residues.

Extending this finding to human MutL homologs, we showed that mutations at the corresponding residues in PMS2 and MLH1—reported as variants of uncertain clinical significance—lead to substantial or complete loss of the ATPase activity. Given that even modest reductions in the MutL ATPase activity are known to severely compromise mismatch repair efficiency *in vivo*, our findings suggest that these variants might be functionally deleterious. Thus, the mechanistic insights obtained in this study provide a structural and biochemical basis for evaluating the pathogenic potential of clinically ambiguous variants in human MutL homologs.

## Materials and Methods

### Construction of expression plasmids

Expression plasmids encoding mutant forms of the aqMutL NTD (residues 1–315), the aqGyrB NTD (residues 1–397), the human PMS2 NTD (residues 1–365), and the human MLH1 NTD (residues 1–337) were generated by introducing point mutations into pET-11a/*aqMutL NTD* ^47^, pET-11a/*aqGyrB NTD* ^11^, pCold ProS2/*human PMS2 NTD* ^35^, and pET28-MLH1/*human MLH1 NTD*^11^ plasmids using the PrimeSTAR site-directed mutagenesis protocol (Takara). Primer sequences used for mutagenesis are listed in Supplementary Table S1. DNA sequencing confirmed that no unintended mutations were introduced.

### Expression and purification of proteins

The wildtype aqMutL NTD, aqGyrB NTD, ProS2-tagged human PMS2 NTD, and His-tagged human MLH1 NTD were overexpressed and purified as described previously ^11,35,47^. The mutant forms were overexpressed and purified using the same procedures as for the respective wildtype proteins.

### CD spectrometry

CD measurements were performed with a spectropolarimeter, model J-720W (Jasco). Measurements of CD spectra were carried out in a solution comprised of 20 mM Tris-HCl (pH 8.0) (eLANT) and 10 μM protein using a 0.1 cm cell at 25°C. The residue molar ellipticity [Θ] was defined as 100 Θ_obs_/(*lc*), where Θ_obs_ is the observed ellipticity, *l* is the length of the light path in centimeters, and *c* is the residue molar concentration of the protein.

### ATPase assay

ATPase activities of the aqMutL NTD, aqGyrB NTD, ProS2-tagged human PMS2 NTD, and His-tagged human MLH1 NTD were measured as described previously ^35^. Reactions were carried out using 10 μM protein in a 20 μl reaction mixture containing 50 mM HEPES–KOH (pH 7.5) (eLANT), 100 mM NaCl, 5 mM MgCl□, and various concentrations of ATP (Roche). For the aqMutL NTD and aqGyrB NTD, reactions were performed at 70°C for 60 min. For the NTDs of the human MutL homologs, reactions were performed at 37°C for 120 min. All experiments were conducted in triplicate. Kinetic parameters were determined by fitting the data to the standard Michaelis–Menten equation using IgorPro.

### Equilibrium dialysis

Equilibrium dialysis experiments were performed as described previously ^11^. Seventy-five microliters of buffer containing 50 mM Tris–HCl (pH 8.0), 100 mM NaCl, 5 mM MgCl□, and 50 μM AMPPNP (Roche) were loaded into the buffer chamber of a DispoEquilibrium Dialyzer equipped with a 10 kDa molecular weight cut-off membrane (Harvard Bioscience, Inc.). An equal volume of the aqMutL NTD (0–320 μM) or aqGyrB NTD (0–400 μM) in the same buffer without AMPPNP was placed in the opposing sample chamber. The device was gently agitated at room temperature for 24 h to allow equilibration. The absorbance of the solution in the buffer chamber was measured at 260 nm. The concentration of unbound AMPPNP was calculated using a molar extinction coefficient (ε = 15,400 M ¹ cm ¹).

### Crystallization and structure determination

Protein crystallization was performed by the sitting-drop vapor diffusion method at 20°C. One microliter of the aqGyrB NTDs (14 mg/ml) containing 0.6 mM AMPPNP (Roche) and 1 mM MgCl was mixed with an equal volume of the reservoir solution and equilibrated against 500 μl of reservoir solution. For the wildtype and D51A mutant forms of the aqGyrB NTD, crystals were obtained using a reservoir solution containing 0.2 M ammonium acetate, 0.1 M HEPES (pH 7.5), and 45% (v/v) 2-methyl-2,4-pentanediol. The E48A mutant form was crystallized in 0.05 M cadmium sulfate hydrate, 0.1 M HEPES (pH 7.5), and 20% (w/v) polyethylene glycol 3,350. The E48Q mutant form was crystallized under identical conditions consisting of 30% (w/v) polyethylene glycol 400, 100 mM HEPES–NaOH (pH 7.5), and 200 mM sodium chloride. The D51N mutant form was crystallized in 0.1 M sodium malonate (pH 7.0) and 12% (w/v) polyethylene glycol 3,350. Crystals of the wildtype and E48A mutant forms were soaked in the respective reservoir solutions supplemented with 30% (v/v) glycerol before being flash-cooled in liquid nitrogen. Crystal of the D51N mutant form was transferred to the reservoir solution containing 30% (w/v) polyethylene glycol 3,350 prior to flash-cooling in liquid nitrogen. Crystal of the D51A mutant form was directly flash-cooled in liquid nitrogen without additional cryoprotection.

X-ray diffraction data were collected at beamline BL45XU in SPring-8 (Hyogo, Japan) at a wavelength of 1.000 Å and a temperature of −173°C. Data were automatically collected and processed using the ZOO system ^48^, in which data processing was assisted by KAMO ^49^. The linearity of the Wilson plot was confirmed over the resolution range employed. Phases were determined by molecular replacement using Phaser with a search model predicted by AlphaFold2 ^50^. Model building and manual correction were performed using COOT ^51^, and structural refinement was carried out with PHENIX refine ^52^. Data collection and refinement statistics are summarized in Table 2. All figures illustrating the protein structures were generated using PyMOL (Schrödinger).

### Phylogenetic analysis and Ancestral state reconstruction

The ATPase domains of representative GHKL superfamily members (MutL, GyrB, MORC, and Hsp90) were aligned using amino acid sequences extracted based on domain boundaries defined by structural correspondence. The multiple sequence alignment was trimmed using trimAl ^53^ to remove ambiguously aligned regions. Maximum-likelihood phylogenetic inference was performed using IQ-TREE (v3.0.1) ^54,55^. Ancestral state reconstruction was performed on the fixed maximum-likelihood topology using IQ-TREE under the empirical Bayesian framework implemented with the selected amino acid substitution model. The state of the amino acid residue at the alignment position corresponding to aqMutL Glu32 or aqGyrB Asp52 was examined. Maximum-likelihood phylogenies were visualized and annotated using the Interactive Tree of Life (iTOL) web server ^56^. Trees were displayed in a rectangular layout without specification of an external outgroup. For visualization purposes, branch lengths were not scaled and were displayed with equal lengths in Fig. 6. The corresponding phylogeny with branch lengths proportional to the inferred evolutionary distances is provided in Supplementary Fig. S5.

## Acknowledgements

The X-ray crystallographic experiments were performed at BL45XU in SPring-8, with the approval of JASRI (proposal nos. 2021A2758, 2021A2758, 2022A2729, and 2022B2729). The authors would like to thank the beamline scientists at SPring-8 BL45XU for their help in collecting the X-ray diffraction data. This work was supported by JSPS KAKENHI Grant Number JP24K08718.

## Author Contributions

KF and TY designed the research; KF, AS, and TM performed experiments; KF, AS, TM, and TY analyzed the data; KF and TY wrote the paper incorporating the opinions of all authors.

## Data Accessibility Statement

The structural data of WT, E48A, E48Q, D51A, and D51N mutant forms of the aqGyrB NTD are available at the protein data bank (https://www2.rcsb.org/) with accession numbers 23UL, 23UV, 23UX, 23UZ, and 23UY, respectively.

## Supplementary Materials

**Supplementary Figure S1.**
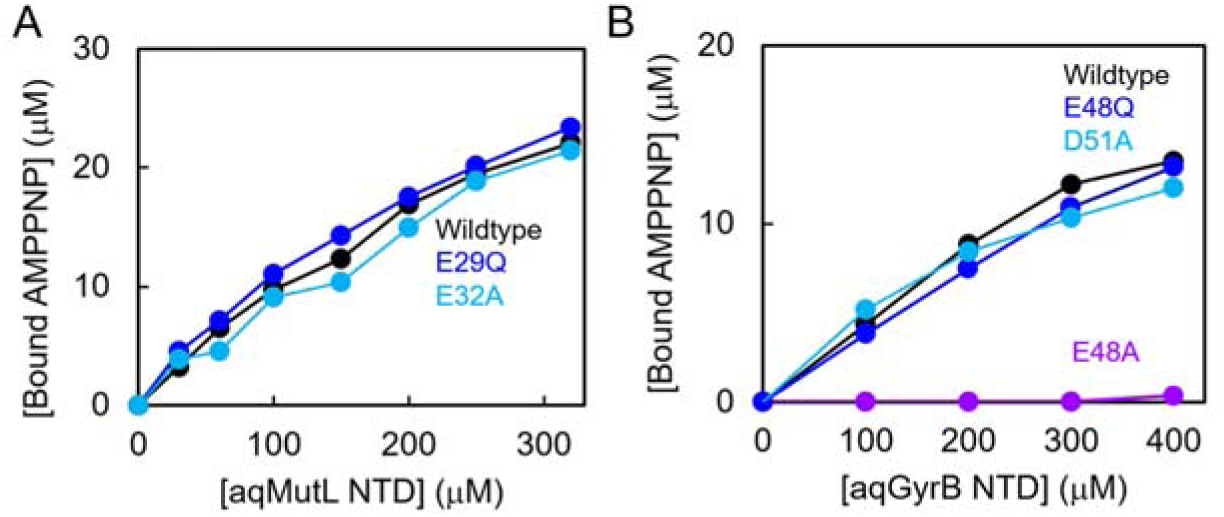
The ATP-binding ability of the aqMutL NTDs (A) and aqGyrB NTDs (B). Binding affinities were quantified by equilibrium dialysis. For each protein concentration, the concentration of unbound AMPPNP was determined from the absorbance at 260 nm of the buffer chamber solution after equilibrium had been reached (see Materials and Methods). The concentration of bound AMPPNP was calculated by subtracting the concentration of unbound AMPPNP from the total AMPPNP concentration and was plotted against the protein concentration.

**Supplementary Figure S2.**
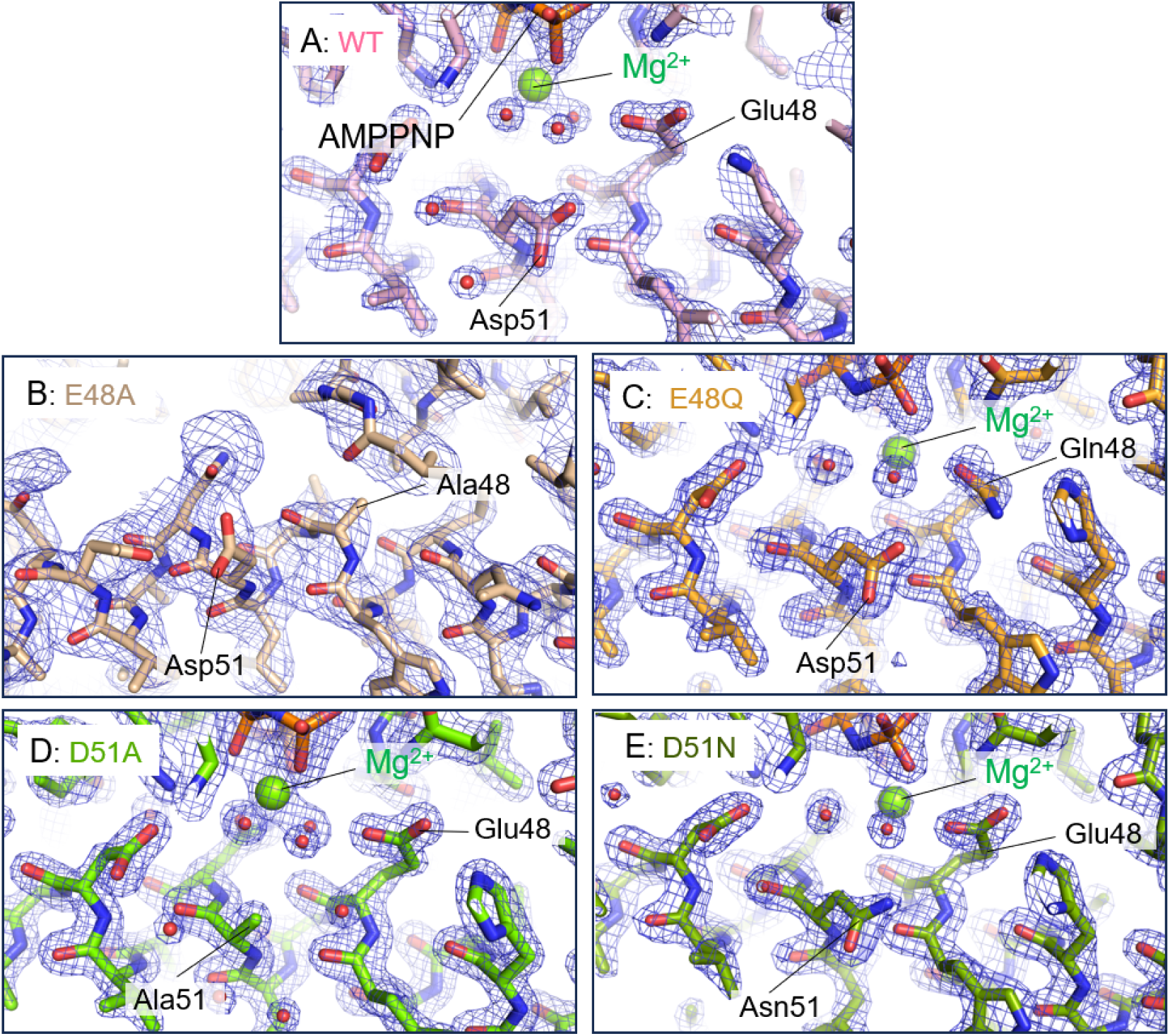
Electron density maps around the catalytic acidic residues in the wildtype and mutant forms of the aqGyrB NTD. Representative 2*F*_o_–*F*_c_ electron density maps (blue mesh, contoured at 3.0σ) around the catalytic acidic residues in the AMPPNP-bound structures of the aqGyrB NTD. (A) Wildtype, (B) E48A, (C) E48Q, (D) D51A, and (E) D51N. The electron density clearly supports the modeled conformations of Glu48, Asp51, and their substituted residues.

**Supplementary Figure S3.**
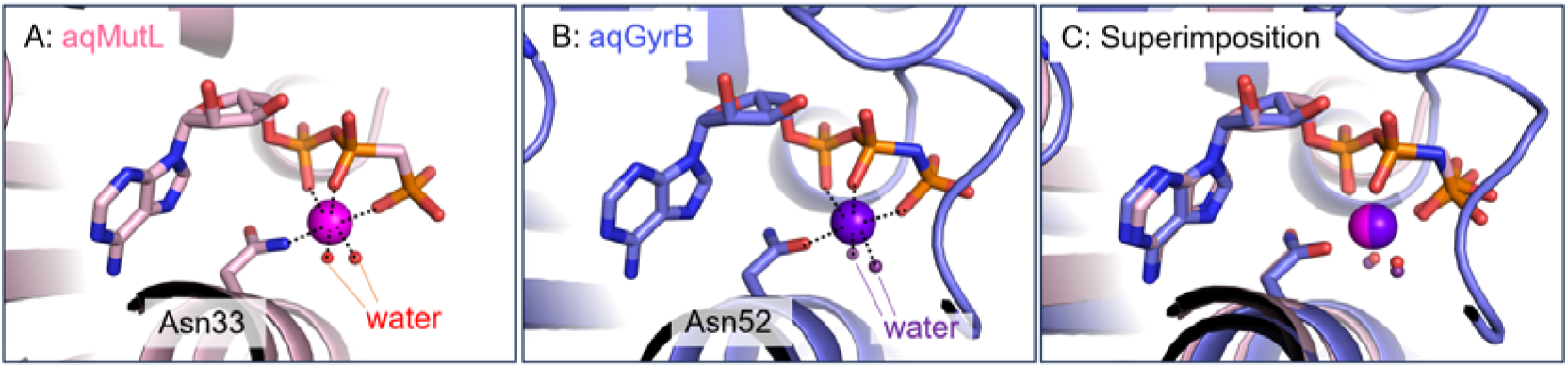
Conserved positioning of the catalytic Mg² ion in the aqMutL NTD and aqGyrB NTD. (A) Close-up view of the ATPase active site of the aqMutL NTD. (B) Close-up view of the ATPase active site of the aqGyrB NTD. (C) Superposition of the active sites of the aqMutL NTD (light pink) and aqGyrB NTD (light blue). Mg² ions are shown as magenta and purple spheres and catalytic water molecules as small spheres. AMPPNP molecules and side chains of Asn33 of the aqMutL NTD and Asn52 of the aqGyrB NTD are shown as stick models. These Mg² -coordinating asparagine residues are adjacent to, but distinct from, the second acidic residues Glu32 in aqMutL and Asp51 in aqGyrB examined in this study. The superposition demonstrates that the catalytic Mg² ion occupies essentially the same position in the aqMutL and aqGyrB active sites.

**Supplementary Figure S4.**
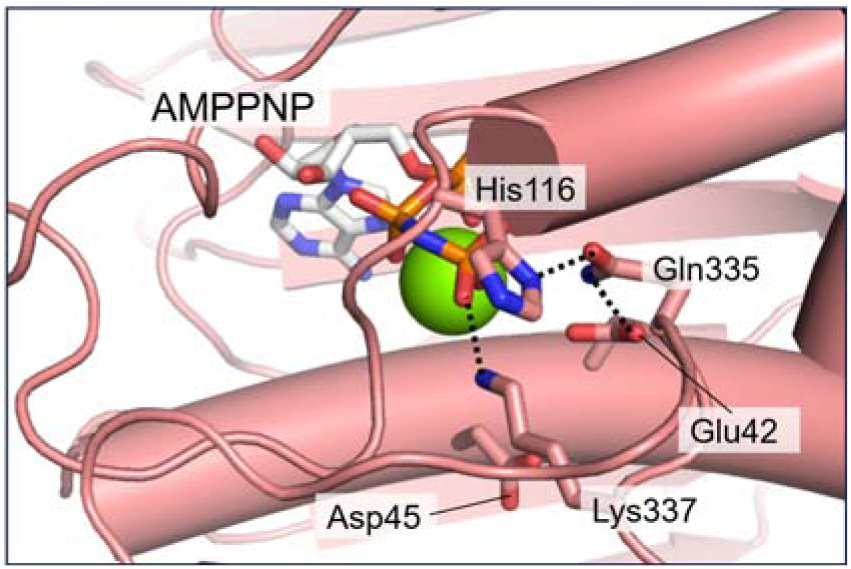
Structure of the ATPase active site of the *E. coli* GyrB NTD. Residues corresponding to Glu48, Asp51, His122, Gln340, and Lys342 of aqGyrB are Glu42, Asp45, His116, Gln335, and Lys337, respectively, in *E. coli* GyrB. Side chains of these residues are shown as stick models together with the bound AMPPNP. Hydrogen bonds and ionic interactions are depicted as dashed lines.

**Supplementary Fig. S5.**
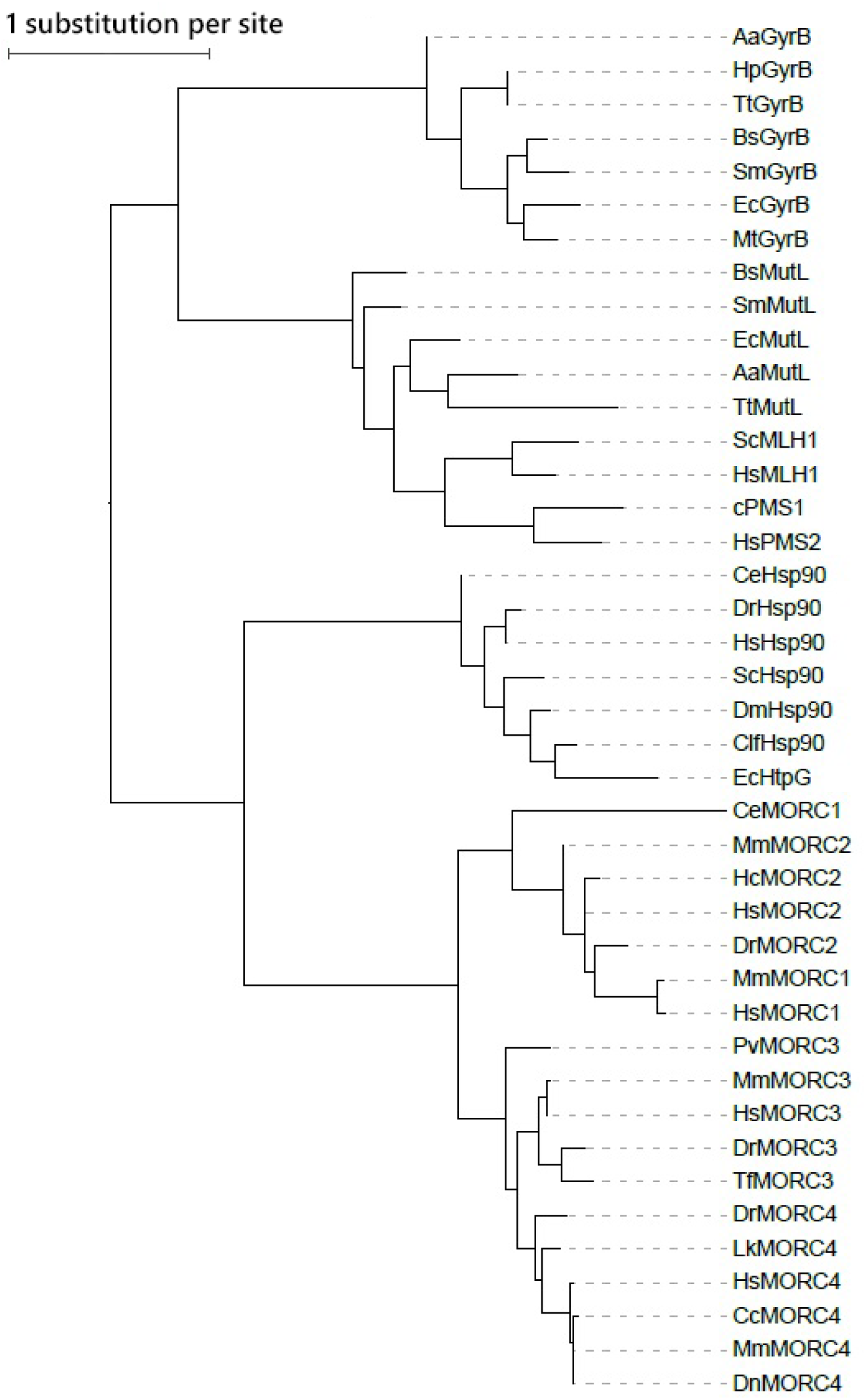
Maximum-likelihood phylogeny corresponding to Fig. 6 with branch lengths proportional to the inferred evolutionary distances. The topology is identical to that shown in Fig. 6, whereas branch lengths are proportional to the evolutionary distances inferred by IQ-TREE. The tree was visualized using the Interactive Tree of Life (iTOL) web server. Branch lengths are proportional to the evolutionary distances inferred by IQ-TREE. The scale bar represents an evolutionary distance of one amino acid substitution per site. Species abbreviations are the same as those in Fig. 6.

**Supplementary Table S1.**
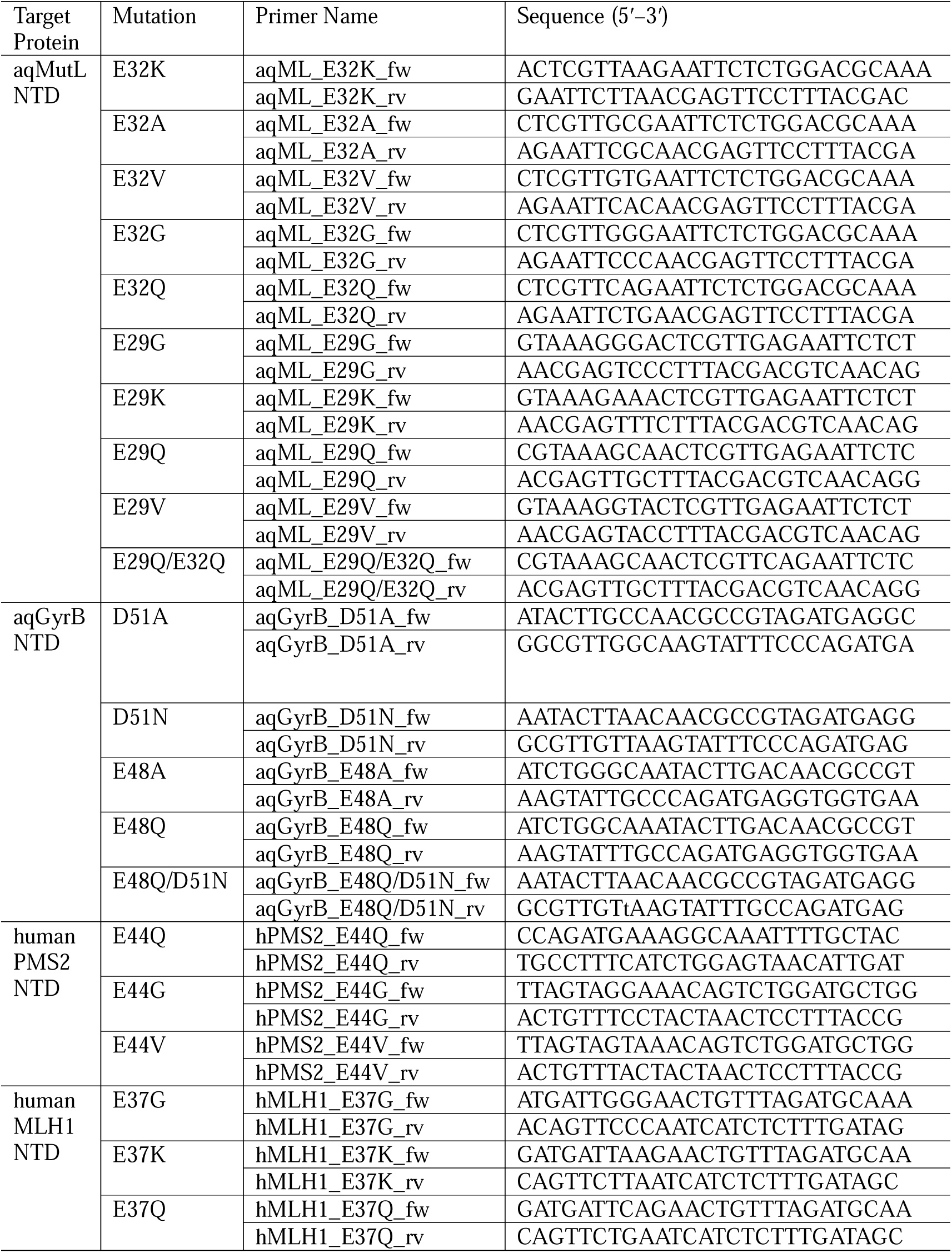
Primers used for site-directed mutagenesis in this study.

